# The demographic and adaptive history of the African green monkey

**DOI:** 10.1101/098947

**Authors:** Susanne P. Pfeifer

**Author notes:** EPFL SV IBI, AAB 048, Station 15, CH-1015 Lausanne, Switzerland.

## Abstract

Relatively little is known about the evolutionary history of the African green monkey (genus *Chlorocebus*) due to the lack of sampled polymorphism data from wild populations. Yet, this characterization of genetic diversity is not only critical for a better understanding of their own history, but also for human biomedical research given that they are one of the most widely used primate models. Here, I analyze the demographic and selective history of the African green monkey, utilizing one of the most comprehensive catalogs of wild genetic diversity to date, consisting of 1,795,643 autosomal single nucleotide polymorphisms in 25 individuals, representing all five major populations: *C. a. aethiops*, *C. a. cynosurus*, *C. a. pygerythrus*, *C. a. sabaeus*, and *C. a tantalus*. Assuming a mutation rate of 5.9 × 10^−9^ per base pair per generation and a generation time of 8.5 years, divergence time estimates range from 523-621kya for the basal split of *C. a. aethiops* from the other four populations. Importantly, the resulting tree characterizing the relationship and split-times between these populations differs significantly from that presented in the original genome paper, owing to their neglect of within-population variation when calculating between population-divergence. In addition, I find that the demographic history of all five populations is well explained by a model of population fragmentation and isolation, rather than novel colonization events. Finally, utilizing these demographic models as a null, I investigate the selective history of the populations, identifying candidate regions potentially related to adaptation in response to pathogen exposure.

## Introduction

The African green monkey (genus *Chlorocebus*), an Old World Monkey also referred to as the vervet monkey, is an abundant primate inhabiting most ecological zones within sub-Saharan Africa (with the exception of tropical forests and deserts (Figure 1; Struhsaker 1967)). Their broad natural geographic distribution, reaching from west to east Africa and from south of the Sahara to the Cape region of South Africa, together with introduced populations on Cape Verde Island and the Caribbean (Groves 2001; Groves 2005), makes vervets an excellent model system to investigate adaptation to local environments. In addition, due to their genetic and physiological similarity to humans, with whom they share a most recent common ancestor roughly 25 million years ago (Mya) (Kumar and Hedges 1998), they are one of the most important non-human primate models in basic and applied biomedical research, widely employed for studies on development, cognition, and behavior (*e.g*., Fairbanks and McGuire 1988; Fairbanks *et al.* 2004; van de Waal and Whiten 2012; Cramer *et al.* 2013), inflammatory, infectious and metabolic disease (Rudel *et al.* 1997; Broussard *et al.* 2001; Olobo *et al.* 2001; Goldstein *et al.* 2006; Ma *et al.* 2014), as well as neurological disorders, in particular Alzheimer's and Parkinson's disease (Lemere *et al.* 2004; Emborg 2007). In contrast to rodents, which are frequently used in biomedical studies but which shared a common ancestor with humans roughly 70Mya, vervet monkeys resemble humans much more closely - not only in their physiology but also in their susceptibility and response to infectious agents, thus making them a particularly valuable model to study pathogens ranging from influenza virus to the simian immunodeficiency virus, a close relative of human immunodeficiency virus.

**Figure 1:**
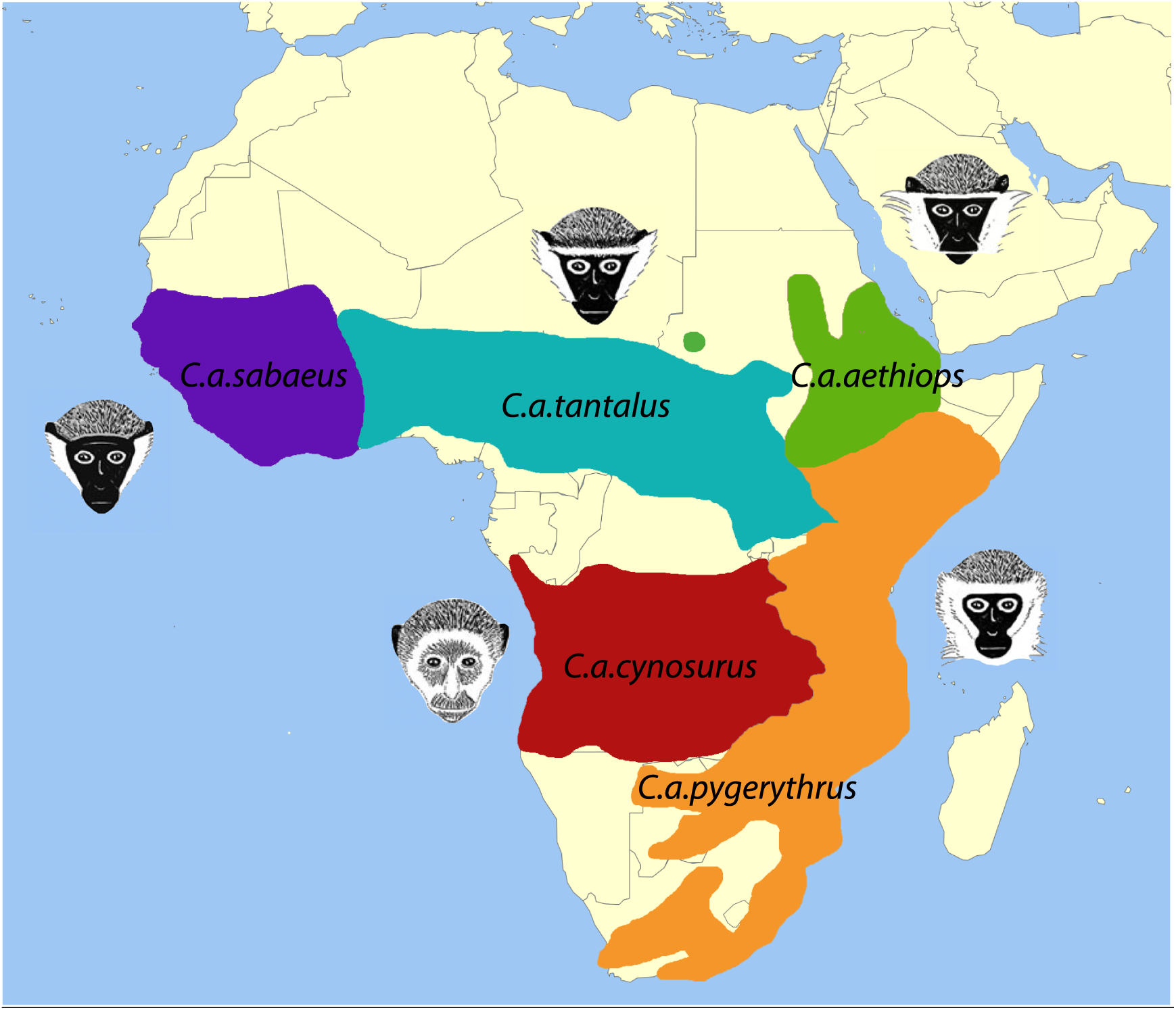
Geographic distribution of the five African vervet monkey populations (red: *C. a. cynosurus;* orange: *C. a. pygerythrus;* green: *C. a. aethiops;* turquise: *C. a. tantalus;* purple: *C. a. sabaeus;* adapted from Haus *et al.* 2013).

Despite the fact that phenotypic and genetic data is publically available from both managed pedigrees and feral populations, relatively little is known about the evolutionary history of the vervet monkey. In fact, the taxonomy of the vervet monkey is disputed and remains the topic of much scientific debate. Groves (2001; 2005) classified vervets into five major species that are phenotypically and geographically distinct: *C. sabaeus* (aka callithrix) inhabiting West Africa (from Senegal to the Volta river), *C. tantalus* (aka tantalus) inhabiting north Central Africa (from Sudan to Ghana and Kenya), *C. pygerythrus* (aka vervet) inhabiting East and Southern Africa (from the eastern Rift Valley in Ethiopia, Somalia, and southern Sudan to South Africa), *C. aethiops* (aka grivet) inhabiting the east of the White Nile region in Ethiopia, as well as areas in Somalia from Khartoum to Mongalla, Eritrea, and Ethiopia, south to the Omo river, and *C. cynosurus* (aka malbrouck) inhabiting Central and Southern Central Africa (from the Albertine Rift to the Atlantic coast as well as northern Namibia and Zambia), with a potentially sixth species, *C. djamdjamensis* (aka the Bale Mountain vervet), limited to small mountain zones in the highland of Ethiopia (Goldstein *et al.* 2006). In contrast, Grubb *et al.* (2003) classified all vervets into a single polytypic species (*Chlorocebus aethiops*). Previous research indicating that vervet monkeys freely interbreed in the narrow contact zones along their geographical boundaries (Detwiler *et al.* 2005; Mekonnen *et al.* 2012; Haus *et al.* 2013) supports the single-species taxonomy, thus I will follow the taxonomy proposed by Grubb *et al.* (2003). Furthermore, the demographic history of the vervet recently inferred using whole genome sequence data of single individuals from each of the major five African populations (Warren *et al.* 2015) is in strong disagreement with earlier work based on smaller data sets and mitochondrial DNA (mtDNA) (Perelman *et al.* 2011; Guschanski *et al.* 2013; Haus *et al.* (2013), regarding both the inferred tree topology as well as the estimated split times.

However, this previous work was severely limited by insufficient polymorphism data sampled from these populations. Fortunately, with the recent availability of a high-quality reference genome (Warren *et al.* 2015), it is now feasible to directly investigate the intra- and interspecific genetic diversity of these different vervet populations, enabling a more accurate view of their demographic history. In this study, whole genome data of five individuals from each of the major five wild African populations was used to infer their demographic history providing much improved clarity to address the conflicting estimates in the literature, and to perform the first naïve scan for genomic targets of positive selection. Thereby, the knowledge gained from better understanding the population genetics of this species may directly benefit medical research in at least two ways. Firstly, by identifying candidate regions under selection, it may be possible to functionally validate phenotype-genotype relationships, particularly with regards to pathogen response. Secondly, understanding the extent of structure present between natural populations is highly important for future genome-wide association studies - as unknowingly sampling across hidden structure can result in spurious results.

## Results and Discussion

Whole genome data of 25 individuals (mean coverage of 4.5X per individual) was used to infer variants and haplotypes in each of the five wild African *Chlorocebus aethiops* populations (see Materials and Methods for details). Across the autosomes, 1,795,643 variants were identified; for which 1,149,007 have an ancestral state that could be unambiguously determined using rhesus macaque as an outgroup. The variants were distributed appropriately across chromosomes (Table 1) and the number of variants identified per sample within a population was highly consistent (Supplementary Figure 1). At regions with sufficient coverage, variation in single nucleotide polymorphism (SNP) density was present, but no strong peaks of SNP density (indicative of a high local false positive rate) were observed. Among the total number of segregating sites in the five populations, 39% were shared between at least two populations and ~1% were shared between all populations, with the remaining polymorphisms being private to a single population (Figure 2).

**Table 1:** Summary of SNP data.

| Chromosome | Length | # SNPs | Ts/Tv | % Accessible | SNPs/kb |
| --- | --- | --- | --- | --- | --- |
| 1 | 126,035,930 | 88,368 | 2.9 | 18.4 | 0.7 |
| 2 | 90,373,283 | 69,432 | 3.1 | 20.0 | 0.8 |
| 3 | 92,142,175 | 61,810 | 2.6 | 16.1 | 0.7 |
| 4 | 91,010,382 | 59,256 | 2.7 | 16.0 | 0.7 |
| 5 | 75,399,963 | 58,652 | 2.9 | 19.2 | 0.8 |
| 6 | 50,890,351 | 35,519 | 3.2 | 16.7 | 0.7 |
| 7 | 135,778,131 | 74,807 | 2.5 | 14.6 | 0.6 |
| 8 | 139,301,422 | 102,193 | 2.8 | 17.7 | 0.7 |
| 9 | 125,710,982 | 92,784 | 2.9 | 18.9 | 0.7 |
| 10 | 128,595,539 | 81,526 | 2.7 | 16.8 | 0.6 |
| 11 | 128,539,186 | 87,591 | 2.8 | 17.0 | 0.7 |
| 12 | 108,555,830 | 74,106 | 2.9 | 18.1 | 0.7 |
| 13 | 98,384,682 | 60,334 | 2.6 | 15.6 | 0.6 |
| 14 | 107,702,431 | 75,455 | 2.9 | 17.9 | 0.7 |
| 15 | 91,754,291 | 59,772 | 2.6 | 16.3 | 0.7 |
| 16 | 75,148,670 | 52,909 | 3.2 | 19.2 | 0.7 |
| 17 | 71,996,105 | 51,115 | 2.7 | 17.2 | 0.7 |
| 18 | 72,318,688 | 54,162 | 2.8 | 17.9 | 0.7 |
| 19 | 33,263,144 | 29,102 | 3.3 | 21.2 | 0.9 |
| 20 | 130,588,469 | 96,660 | 3.0 | 18.8 | 0.7 |
| 21 | 127,223,203 | 83,854 | 2.7 | 16.8 | 0.7 |
| 22 | 101,219,884 | 68,197 | 2.8 | 17.6 | 0.7 |
| 23 | 82,825,804 | 56,430 | 2.8 | 17.7 | 0.7 |
| 24 | 84,932,903 | 55,184 | 2.8 | 16.6 | 0.7 |
| 25 | 85,787,240 | 57,881 | 2.8 | 17.7 | 0.7 |
| 26 | 58,131,712 | 41,674 | 2.9 | 18.8 | 0.7 |
| 27 | 48,547,382 | 33,665 | 2.8 | 17.1 | 0.7 |
| 28 | 21,531,802 | 14,786 | 3.3 | 16.5 | 0.7 |
| 29 | 24,206,276 | 18,419 | 2.8 | 18.0 | 0.8 |
| total | 2,607,895,860 | 1,795,643 | 2.8 | 17.5 | 0.7 |

**Figure 2:**
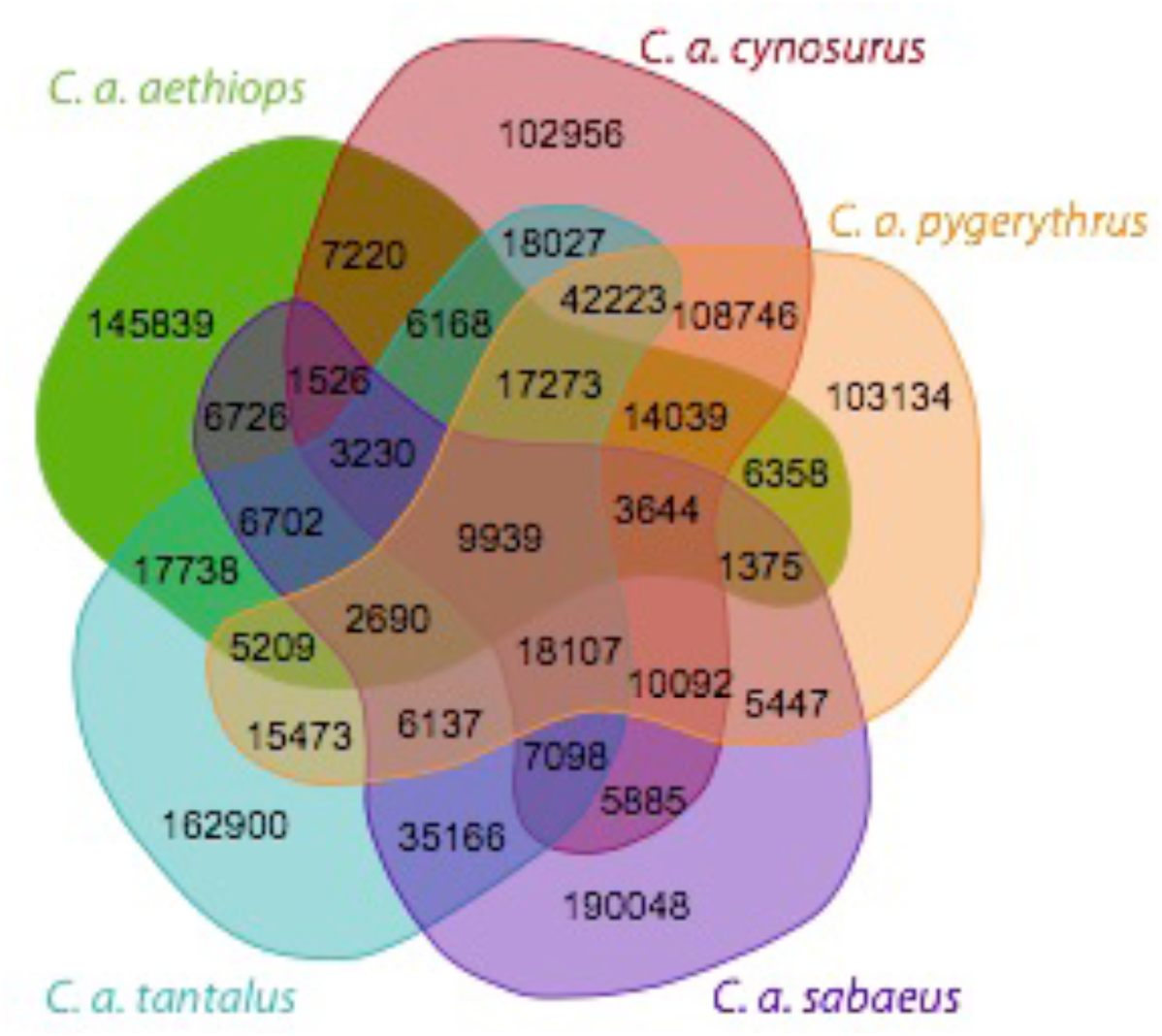
Private and shared segregating sites per population (red: *C. a. cynosurus*; orange: *C. a. pygerythrus*; green: *C. a. aethiops*; turquise: *C. a. tantalus*; purple: *C. a. sabaeus*). Note that the sizes of the areas are not proportional to the magnitude of the numbers.

### Population Structure

The five populations exhibit similar levels of nucleotide diversity (*π*_intergenic_ = 3.7-5.2 × 10^−4^) in the non-coding, non-repetitive parts of the genome (Table 2) - on the lower end of nucleotide diversity levels previously reported in other primates (Yu *et al.* 2004; Fischer *et al.* 2006; Prado-Martinez *et al.* 2013). Nucleotide diversity levels, and thus inferred effective population sizes (*N_e_*), are highest in *C. a. cynosurus* and *C. a. tantalus,* and lowest in *C. a. aethiops,* with intermediate levels in *C. a. pygerythrus* and *C. a. sabaeus*. The level of genetic differentiation between populations (as assessed by weighted *F_st_*), ranging from 0.33 (C. *a. cynosurus* - *C. a. tantalus*) to 0.6 (*C. a. aethiops* - *C. a. sabaeus*) (Table 2), indicates strong genetic structure between different populations, overlapping the range of differentiation previously reported between different chimpanzee populations (*e.g., F_st_*(western chimpanzee - central chimpanzee) = 0.25-0.38, and F*st*(western chimpanzee - eastern chimpanzee) = 0.31-0.42; Becquet *et al.* 2007; Fischer *et al.* 2011) as well as bonobos and chimpanzees (*e.g.,* Fst(bonobo - central chimpanzee) = 0.49-0.54 and Fst(bonobo - western chimpanzee) = 0.68-0.74; Fischer *et al.* 2006, 2011). The only exception is a weighted *F_st_* of 0.16 between *C. a. cynosurus* and *C. a. pygerythrus*, similar to *F_st_* values observed between human populations (Rosenberg *et al.* 2002). The large differentiation of *C. a. aethiops* compared with the other populations provides the first line of evidence suggesting that they may represent the earliest split on the tree, contrary to previous results.

**Table 2:** Summaries of genetic variation.

|  | *C. a. aethiops* | *C. a. cynosurus* | *C. a. pygerythrus* | *C. a. sabaeus* | *C. a. tantalus* |
| --- | --- | --- | --- | --- | --- |
| $\pi_{\text{intergenic}}$ | $3.7 \times 10^{-4}$ | $5.2 \times 10^{-4}$ | $5.1 \times 10^{-4}$ | $4.7 \times 10^{-4}$ | $5.2 \times 10^{-4}$ |
| $\pi_{\text{exonic}}$ | $1.4 \times 10^{-4}$ | $2.1 \times 10^{-4}$ | $2.1 \times 10^{-4}$ | $1.9 \times 10^{-4}$ | $1.9 \times 10^{-4}$ |
| $\theta_{W; \text{intergenic}}$ | $3.2 \times 10^{-4}$ | $4.7 \times 10^{-4}$ | $4.6 \times 10^{-4}$ | $4.3 \times 10^{-4}$ | $4.7 \times 10^{-4}$ |
| $\theta_{W; \text{exonic}}$ | $1.3 \times 10^{-4}$ | $1.9 \times 10^{-4}$ | $1.9 \times 10^{-4}$ | $1.7 \times 10^{-4}$ | $1.8 \times 10^{-4}$ |
| $N_e$ | 15,849 | 21,921 | 21,733 | 19,824 | 22,146 |
| $D_{\text{intergenic}}$ | 0.27 | 0.21 | 0.23 | 0.19 | 0.23 |
| $D_{\text{exonic}}$ | 0.06 | 0.06 | 0.07 | 0.06 | 0.06 |
| $F_{st}(\text{aet})$ | - | 0.55 | 0.56 | 0.60 | 0.53 |
| $F_{st}(\text{cyn})$ | 0.55 | - | 0.16 | 0.45 | 0.33 |
| $F_{st}(\text{pyg})$ | 0.56 | 0.16 | - | 0.46 | 0.35 |
| $F_{st}(\text{sab})$ | 0.60 | 0.45 | 0.46 | - | 0.40 |
| $F_{st}(\text{tan})$ | 0.53 | 0.33 | 0.35 | 0.40 | - |

Nucleotide diversity $\pi$ (Nei and Li, 1979) and Watterson's estimate of $\theta$ , $\theta_W$ , (Watterson 1975) were calculated in the non-coding, non-repetitive ( $\pi_{\text{intergenic}}$ ; $\theta_{W; \text{intergenic}}$ ) and exonic ( $\pi_{\text{exonic}}$ ; $\theta_{W; \text{exonic}}$ ) parts of the genome for each population (aet: *C. a.* *aethiops*; cyn: *C. a. cynosurus*; pyg: *C. a. pygerythrus*; sab: *C. a. sabaeus*; tan: *C. a. tantalus*) using the libsequence package msstats v.0.3.4 (Thornton 2003). Effective population sizes ( $N_e$ ) were estimated from the data by fixing the mutation rate $\mu$ to $5.9 \times 10^{-9}$ per base pair per generation (*i.e.*, the mutation rate observed in rhesus macaque as there is no direct estimate for $\mu$ available in vervet monkeys; Hernandez *et al.* 2007). Using VCFtools v.0.1.12b (Danecek *et al.* 2011), Tajima's $D$ (Tajima 1989) was calculated (in 10kb windows along the genome) to test for deviations from the neutral equilibrium site frequency distribution and Weir and Cockerham's fixation index $F_{st}$ was calculated between each pair of populations to assess population differentiation.

To explore genetic evidence for population structure among vervets, the level of shared ancestry among individuals was studied using independent-loci admixture models as implemented in the software fastSTRUCTURE (Raj *et al.* 2014). fastSTRUCTURE assigns individuals to a hypothesized number of populations such that the amount of linkage disequilibrium across loci is minimized (see Materials and Methods). The best-fit model had four ancestry components, strongly supporting the division of samples into four discontinuous populations. The inferred clusters correspond well to the reported labels of *C. a. aethiops*, *C. a. sabaeus*, *C. a. tantalus,* grouping *C. a. cynosurus* and *C. a. pygerythrus* into the same cluster, with little evidence for admixture between the populations (Supplementary Figure 2). fastSTRUCTURE also has power to assess ancestry proportions for individuals with mixed ancestry, of which none were identified. In addition, principal component analysis (PCA) was used to determine the levels of genetic differentiation between individuals. The first principal component, explaining 17.4% of the variation, clearly separates the geographically most isolated population with the smallest habitat range, *C. a. aethiops*, from the other four populations (Figure 3). *C. a. sabaeus*, *C. a. tantalus*, *C. a. cynosurus*, and *C. a. pygerythrus* are distributed along the second principal component (explaining 14.5% of the variation), with no clear differentiation between *C. a. cynosurus* and *C. a. pygerythrus*, both of which inhabit central and south Africa. Construction of an autosomal consensus tree (*i.e.,* a tree that most commonly represents the relationship between the sample locations) indicates that one *C. a. pygerythrus* individual (SRR556103) falls within the *C. a. cynosurus* cluster, whereas separate monophyletic groups are supported for *C. a. aethiops*, *C. a. sabaeus*, and *C. a. tantalus* (Supplementary Figure 3) - consistent with the results of both the fastSTRUCTURE and PCA analyses, and additionally supported by the identity-by-state pattern observed for each pair of individuals (Supplementary Figure 4).

**Figure 3:**
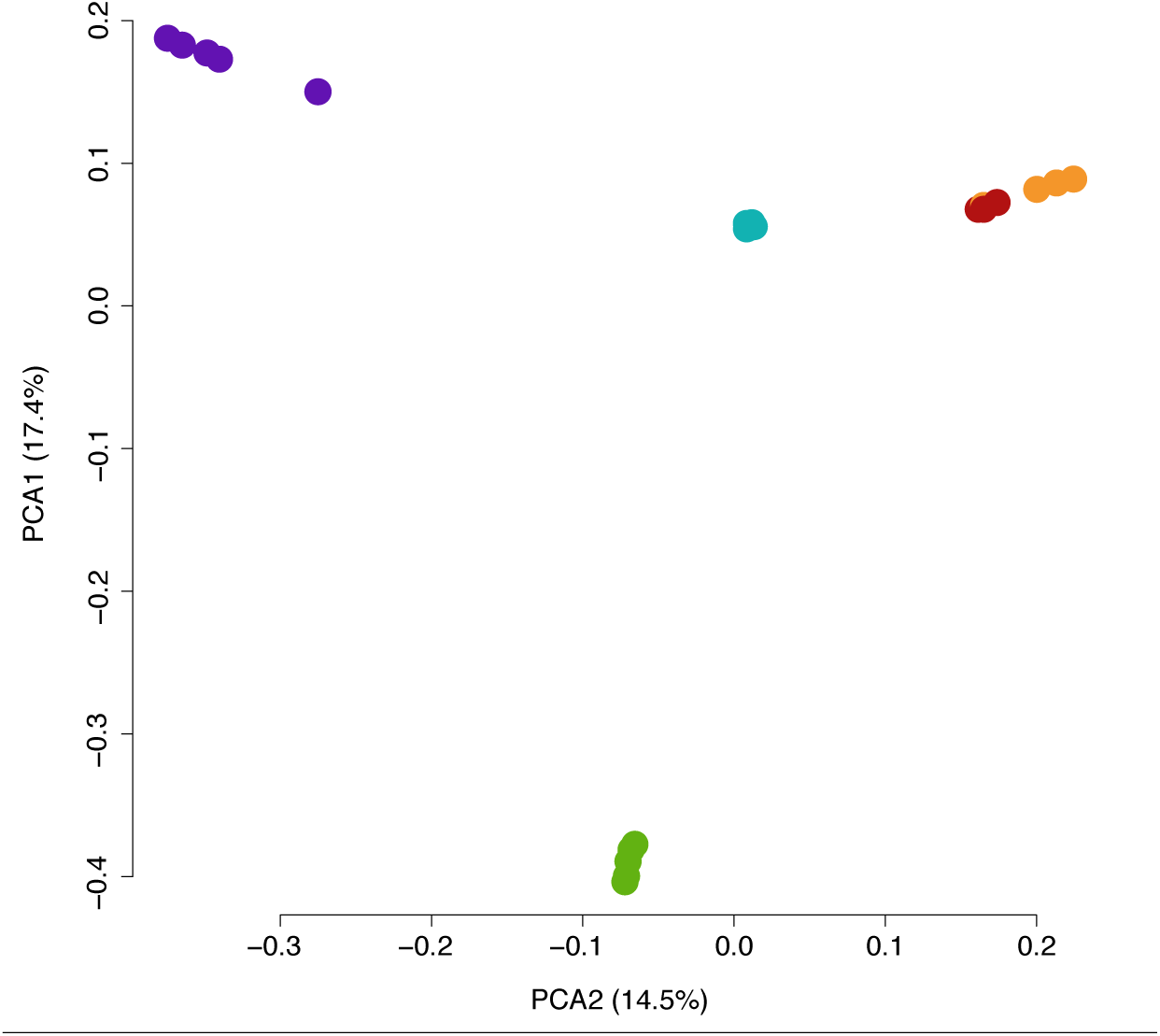
Principal component analysis of all sampled individuals in the different populations (red: *C. a. cynosurus*; orange: *C. a. pygerythrus*; green: *C. a. aethiops*; turquise: *C. a. tantalus*; purple: *C. a. sabaeus*). Data was thinned to exclude SNPs with an r^2^ > 0.2 in order to avoid a strong influence of SNP clusters in the PCA. Percentages indicate the variance explain by each principle component.

### Demographic history

Patterns of genetic divergence were used to elucidate the evolutionary history of the vervet populations. Specifically, divergence times were estimated using a molecular clock based on putatively neutral fixed differences between the genomes of the populations (Table 3), assuming that mutations occurred at a constant rate of 5.9 × 10^−9^ per base pair per generation among lineages (Hernandez *et al.* 2007) and a generation time of 8.5 years (Warren *et al.* 2015). The inferred topology (Figure 4) is consistent with the observed levels of genetic differentiation (Table 2) as well as allele sharing between the species, with evidence of incomplete lineage sorting being observed (Figure 2), as reported for many other primates (Patterson *et al.* 2006; Hobolth *et al.* 2007). The basal split of *C. a. aethiops* from the other four populations was estimated to be 523-621kya. The time estimates for (*C. a. sabaeus* + *C. a tantalus*)/(*C. a. cynosurus* + *C. a. pygerythrus*), *C. a. sabaeus*/*C. a tantalus*, and *C. a. cynosurus*/*C. a. pygerythrus* were 242-333kya, 239kya, and 143kya, respectively. These estimated divergence times are roughly consistent with the ones recently inferred using whole genome sequence data of single individuals from each of the five African populations (Warren *et al.* 2015) but considerably younger than those reported from mtDNA (Guschanski *et al.* 2013). However, the inferred tree topology is in disagreement with results of previously published studies on the topic (Perelman *et al.* 2011; Guschanski *et al.* 2013; Haus *et al.* 2013; Warren *et al.* 2015). Discrepancies in both divergence times and tree shapes between estimates obtained from mitochondrial and nuclear genomic data have been observed in many different species (Avise 1994; Funk and Omland 2003; Chan and Levin 2005; Toews and Brelsford 2012) including other primates (*e.g.,* Wise *et al.* 1997; Stone *et al.* 2010; Nietlisbach *et al.* 2012) - a discordance that has generally been attributed to differences in the selective and demographic histories of mitochondrial and nuclear DNA (such as sex-biased dispersal, different effective female than male population sizes, and adaptive introgression (Toews and Brelsford 2012)), as well as incomplete lineage sorting. The fourfold smaller effective population size in mtDNA compared to nuclear DNA will also make it more susceptible to stochastic variation, thus mtDNA may not be a good representative of ancestry and genetic diversity across the entire genome. In contrast, the discrepancy in tree topology with the one recently inferred by Warren *et al.* (2015) using whole genome sequence data of single individuals from each of the major five African populations is likely explained by the fact that the authors were unable to discern between segregating sites and fixed differences in the populations, owing to the analysis of single individuals. That is, within-population variation was confounded with between-population divergence. In addition, their analyses were based on genome-wide variant data, potentially biasing demographic inferences via both direct and linked selection (see Ewing and Jensen 2016), whereas I here define putatively neutral regions for such analysis.

**Table 3:** Putatively neutral fixed differences between the populations.

|  | *C. a. aethiops* | *C. a. cynosurus* | *C. a. pygerythrus* | *C. a. sabaeus* | *C. a. tantalus* |
| --- | --- | --- | --- | --- | --- |
| *C. a. aethiops* | - | 52,841 | 53,239 | 62,716 | 53,180 |
| *C. a. cynosurus* | 52,841 | - | 14,458 | 32,849 | 24,427 |
| *C. a. pygerythrus* | 53,239 | 14,458 | - | 33,631 | 25,241 |
| *C. a. sabaeus* | 62,716 | 32,849 | 33,631 | - | 24,098 |
| *C. a. tantalus* | 53,180 | 24,427 | 25,241 | 24,098 | - |

**Figure 4:**
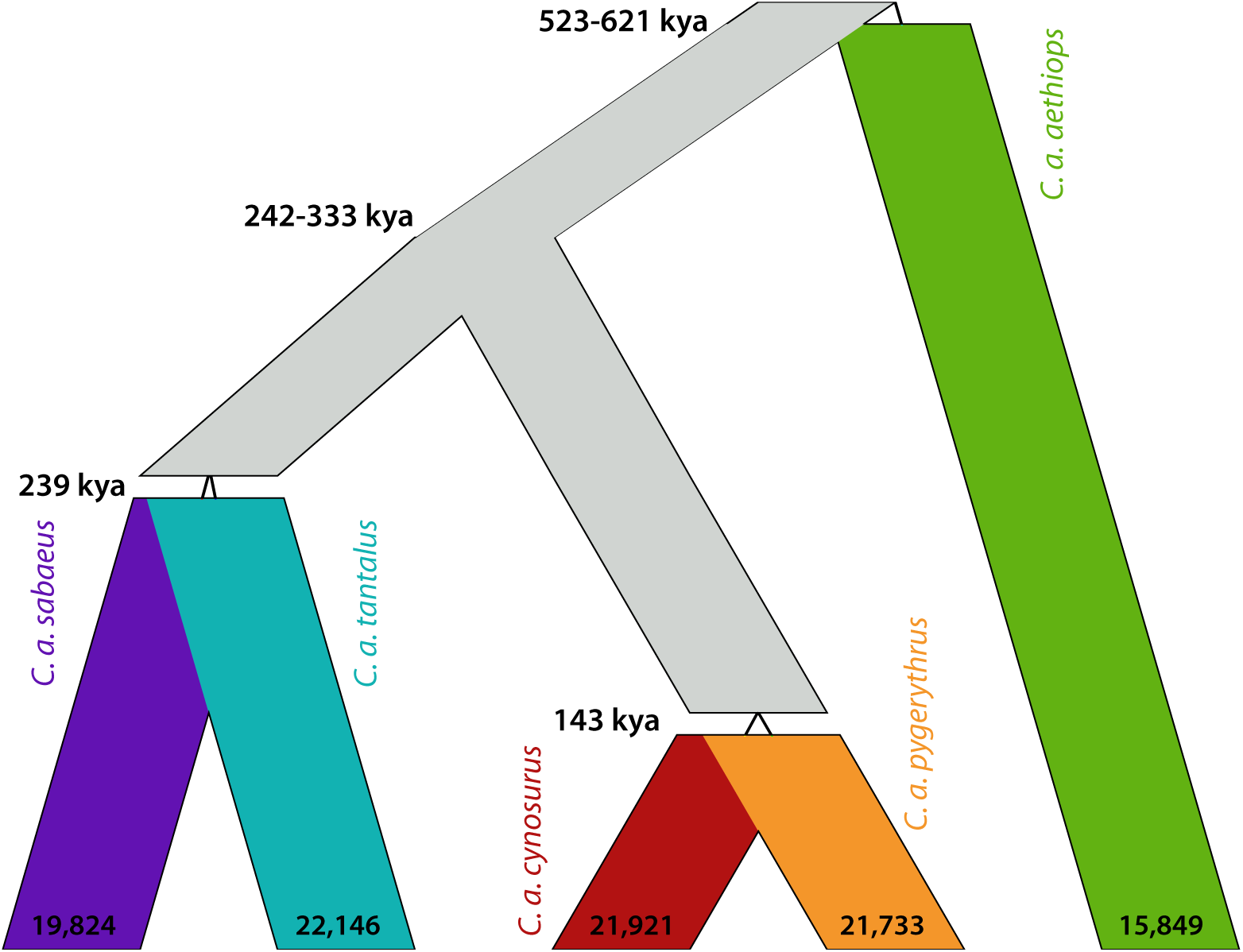
Demographic history of the vervet monkey. Divergence times have been estimated using a molecular clock based on putatively neutral, fixed differences between the genomes of the populations (Table 3), assuming that mutations occurred at a constant rate of 5.9 × 10^−9^ per base pair per generation among lineages (Hernandez *et al.* 2007) and that the generation time is 8.5 years (Warren *et al.* 2015). Effective population sizes (provided at the tip of the branches) were estimated from the data by fixing the mutation rate *μ* to 5.9 × 10^−9^ per base pair per generation (Hernandez *et al.* 2007). The figure was generated using PopPlanner (Ewing *et al.* 2015).

A standard equilibrium model without migration was the best fit to the data (Figure 5), suggesting population fragmentation rather than colonization as a driver of the demographic history of vervet monkeys, as well as historically stable population sizes. Consistent with biases expected from utilizing a multi-sample genotype calling strategy on low coverage sequencing data (Han *et al.* 2013), rare variants were under-called in the data set, distorting the site frequency spectrum (SFS) to include fewer singletons than doubletons. This deficit of low frequency variants results in a stronger underestimation of *π* compared to *θ_W_*, skewing Tajima’s *D* towards slightly positive values in all populations (Table 2). Importantly however, Han *et al.* (2013) demonstrated that rank-based statistics used for genome-wide selection scans are less sensitive to such biases in the inferred SFS, enabling identification even at low coverage.

**Figure 5:**
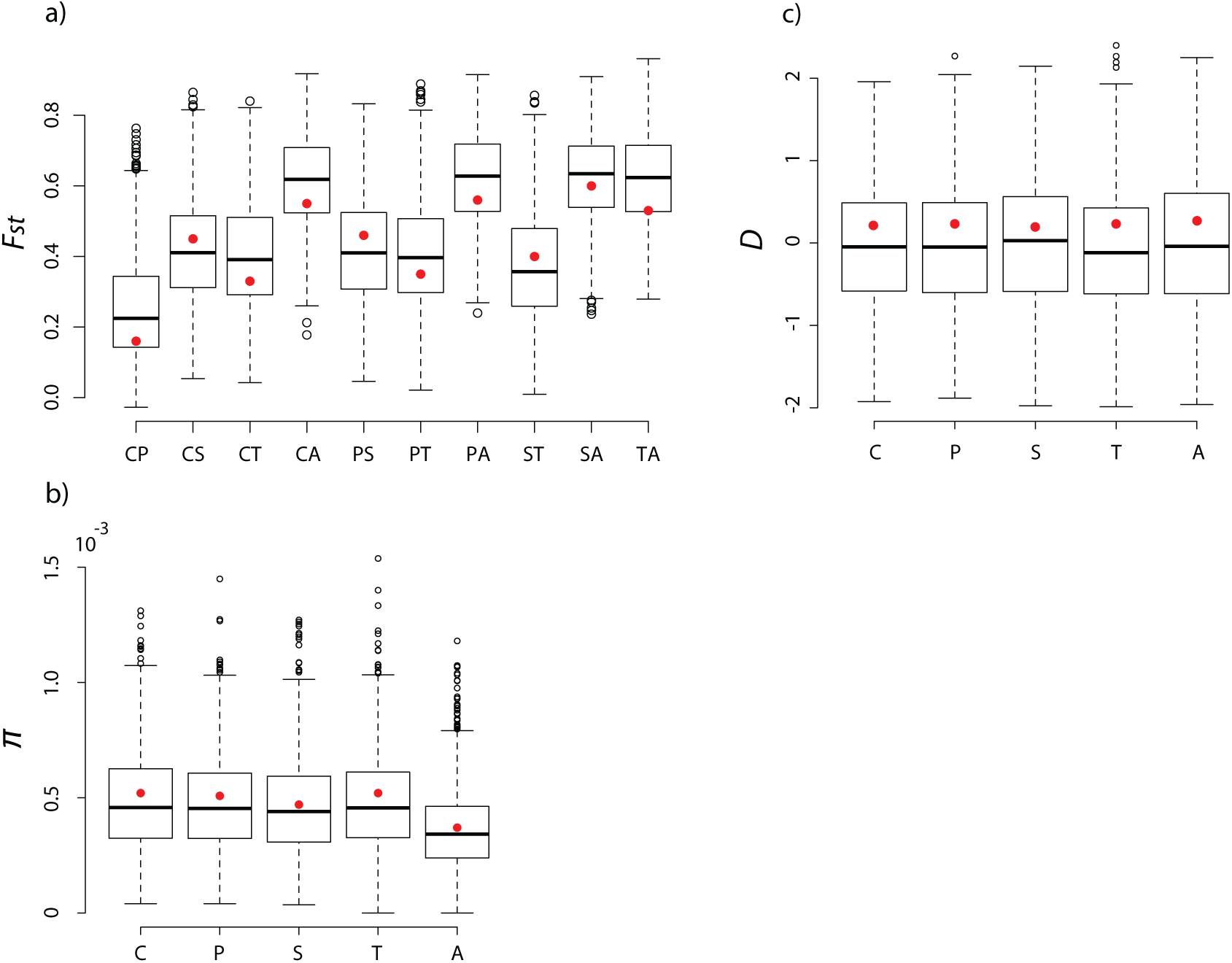
Fit of the data to a standard neutral equilibrium model, using the described putatively neutral class of SNPs. a) Levels of genetic differentiation between populations (A: *C. a. aethiops*; C: *C. a. cynosurus*; P: *C. a. pygerythrus*; S: *C. a. sabaeus*; T: *C. a. tantalus*) observed in the data (red dots) are expected under this model (simulation results shown as boxplots). b) Levels of nucleotide diversity in the vervet monkey populations. c) Tajima’s *D* is skewed towards positive values in all populations compared to the standard equilibrium model due to a deficit of low frequency variants in the data, which is an artifact from utilizing a multi-sample genotype calling strategy on low coverage sequencing data (Han *et al.* 2013).

### Selection

Given the already large divergence between populations shown above, *F_st_*–based scans will be underpowered. Correspondingly, no significant outliers were detected when such an approach was applied to the data (BayeScan; Foll and Gaggiotti 2008). Thus, population-specific scans relying on site frequency spectrum–based expectations were utilized. Statistical tests based on a classical selective sweep model (SweeD; Pavlidis *et al.* 2013) suggest that 1.5-3.1% of the vervet genome may be affected by recent selective sweeps (Supplementary Figure 5), when assuming a 1% false positive rate. Comparisons with the extent of positive selection in other primate genomes are somewhat tenuous, given the strongly differing results depending on the methodology employed (see discussion in Biswas and Akey 2006; Crisci *et al.* 2012; Jensen 2014). It should further be noted that the false discovery rate of selection scans may be higher than anticipated (Teshima *et al.* 2006), due to the challenges in detecting the footprint of a selective sweep. In particular, differentiating sweeps from other patterns of genetic background variation that reflect unaccounted population history, variability in the underlying mutation and recombination landscapes, as well as differing modes of selection, has been challenging (see reviews of Bank *et al.* 2014; Jensen *et al.* 2016). Fortunately, the demographic inference above suggests relatively stable population histories, devoid of the kind of severe population size changes which have been shown to induce major difficulties when conducting genomic scans (Thornton and Jensen 2007). In addition, several regions have strong and consistent evidence of being targeted by positive selection in multiple populations (Supplementary Table), a particularly promising result given both the old split times as well as the lacking evidence for on-going migration. One such region with the strong support includes the gene *DYNC1I1* (Figure 6), a known target of herpes simplex virus, which interacts with the dynein motor of the nuclear membrane to transport capsid-tegument structures to the nuclear pore (Ye *et al.* 2000). This finding is noteworthy given the well-described infection history of vervet monkeys with the herpes virus (Malherbe and Harwin 1958; Clarkson *et al.* 1967; Wertheim *et al.* 2014). Further, this offers an excellent candidate region for investigating host-shift between humans (in which infection is associated with severe symptoms, and may be fatal) and vervets (in which infections are generally asymptomatic) (Nsabimana *et al.* 2008).

**Figure 6:**
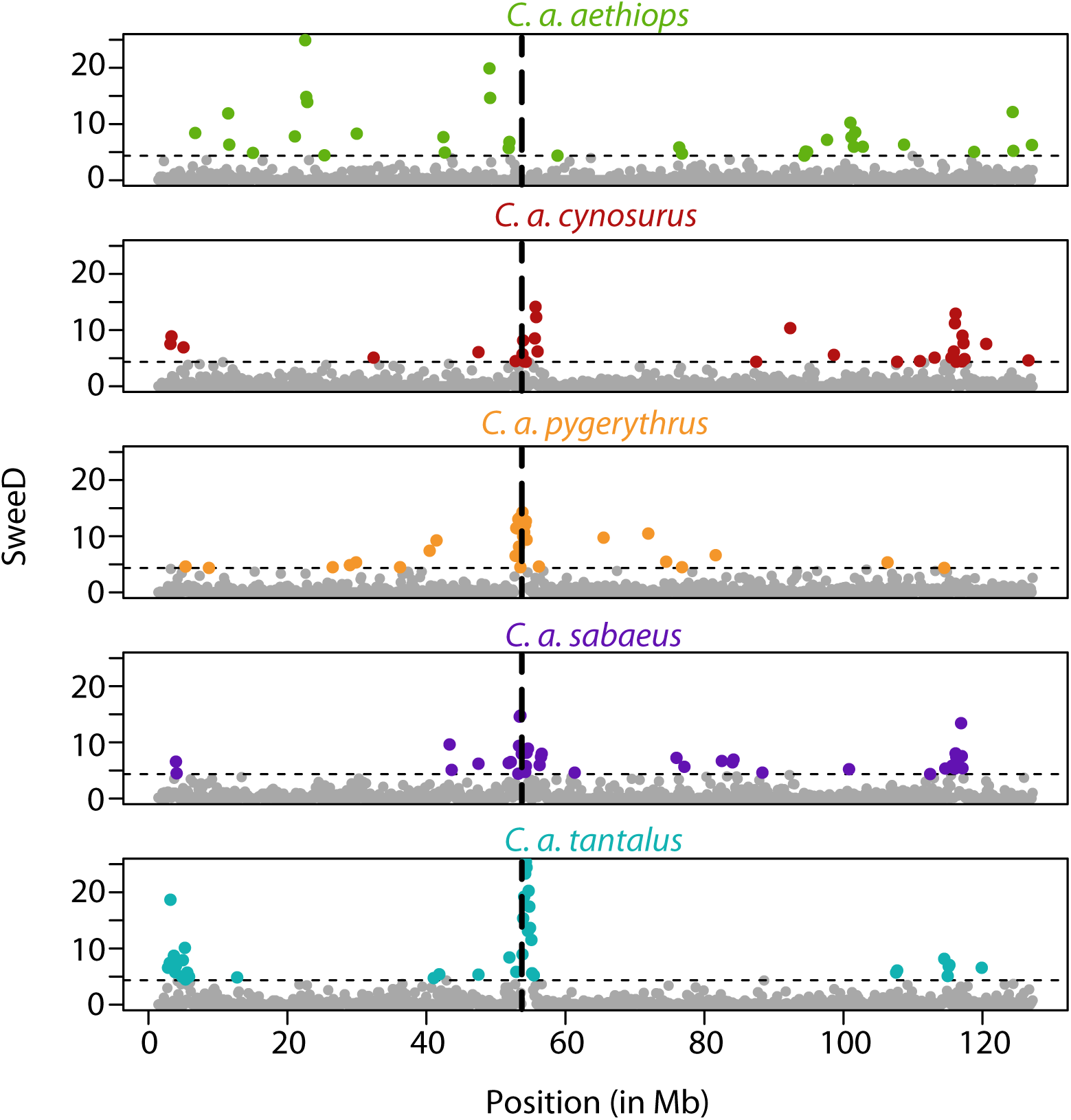
Likelihood surfaces of the CLR test calculated by SweeD (Pavlidis *et al.* 2013) for chromosome 21 per population (green: *C. a. aethiops*; red: *C. a. cynosurus*; orange: *C. a. pygerythrus*; purple: *C. a. sabaeus*; turquise: *C. a. tantalus*) on a megabase (Mb) scale. The dashed black horizontal line is the significance threshold of the test (based on a standard equilibrium model without migration; see Materials and Methods). A selective sweep near the gene *DYNC1I1* (dashed black vertical line; SweeD-score: >14.8) is common to multiple populations.

### Conclusions

To clarify the existing and conflicting estimates regarding the demographic history of the vervet monkey and to characterize the species’ adaptive history, one of the most comprehensive catalogs of wild genetic diversity to date was generated, consisting of 1,795,643 autosomal single nucleotide polymorphisms, identified in five individuals from each of the major five wild African populations. Population genetic analyses confirmed strong genetic structure between the different populations, with nucleotide diversity levels on the lower end of previously reported values in other primates. Divergence between *C. a. aethiops* and the other four extant populations is best fit with a model of population fragmentation and isolation, rather than novel colonization events, beginning roughly 523-621kya. This finding is in strong disagreement with previously published results based on smaller data sets, single individuals, and mitochondrial DNA (Perelman *et al.* 2011; Guschanski *et al.* 2013; Haus *et al.* 2013; Warren *et al.* 2015). The ability to here utilize polymorphism data for improved demographic inference, to account for segregating variation when inferring divergence times, and to focus on putatively neutral regions of the genome, have all contributed to this improved clarity. Further, evidence of recent selective sweeps at the genomic level was detected in all populations. While population-specific events are of interest, the most promising candidates are likely those with evidence in multiple populations. The strongest such signal contains a gene related to viral exposure, providing a valuable candidate for future study concerning both host-shift and the underlying causes of symptomatic infections of the herpes simplex virus.

## Materials and Methods

### Read Mapping

Whole-genome sequence data for 25 individuals (five individuals per population with a genome-wide mean coverage of 4.5X per individual) was downloaded from SRA (*i.e., C. a. tantalus:* SRR556154, SRR556127, SRR556105, SRR556122, SRR556151; *C. a. sabaeus:* SRR556189, SRR556192, SRR556194, SRR556191, SRR556193; *C. a. pygerythrus:* SRR556143, SRR556109, SRR556118, SRR556116, SRR556103; *C. a. cynosurus:* SRR556153, SRR556149, SRR556117, SRR556164, SRR556161; *C. a. aethiops:* SRR556111, SRR556121, SRR556162, SRR556165, SRR556133). Reads were aligned to the repeat-masked *Chlorocebus sabaeus* reference assembly v.1.1 (NCBI GenBank accession number GCA_000409795.2), consisting of assemblies for 29 autosomes (mean length: 89,933,368bp), chromosome X (130,038,232bp), chromosome Y (6,181,219bp), the mitochondrion (16,550bp), as well as 1,972 unplaced scaffolds (mean length: 23,085bp) (Warren *et al.* 2015), and the Epstein-Barr virus genome (NCBI Reference Sequence NC_007605.1) using BWA-MEM v.0.7.13 (Li and Durbin 2009). Thereby, the Epstein-Barr virus assembly was included as a decoy in the read alignment step to enable the absorption of reads that did not originate from vervet monkey DNA (as DNA sequences of interest are often contaminated (*e.g.,* by Epstein-Barr virus, frequently used in laboratories to immortalize the cell lines)). The inclusion of such a decoy genome has been shown to frequently improve the accuracy of read alignments by reducing false positive variant calls (see review of Pfeifer (2016)). Following the Genome Analysis Toolkit (GATK) v.3.5 Best Practice (McKenna *et al.* 2010; DePristo *et al.* 2011; Van der Auwera *et al.* 2013), duplicates were marked using Picard v.2.1.1, before conducting multiple sequence realignments with simultaneous adjustment of Base Alignment Qualities (Li 2011). Next, base quality scores were recalibrated using GATK's BaseRecalibrator v.3.5 together with ~500k variants from the genome-wide SNP panel of the Vervet Genetic Mapping Project (Huang *et al.* 2015), downloaded from the European Variant Archive (study number PRJEB7923).

### Variant Calling and Filtering

Variants were called using GATK’s HaplotypeCaller v.3.5, a method well suited for low coverage depths averaging 4-6X per individual (Cheng *et al.* 2014), and jointly genotyped using GATK’s GenotypeGVCFs v.3.5. Although soft filtering methods using machine learning (such as GATK’s VQSR) have a better specificity at low coverage than hard filtering methods (Cheng *et al.* 2014), these techniques can not readily be applied in this study due to the fact that soft filtering methods require the construction of a statistical model based on a set of known high-quality variant calls. Hence, these methods require a large training data set of known high-quality variants in the underlying model, which unfortunately is not yet available for vervet monkeys. Thus, post-genotyping, the raw variant call set was limited to autosomal, bi-allelic single nucleotide polymorphisms (SNPs) and conservatively filtered using the following hard filtering criteria (with acronyms as defined by the GATK package), attempting to minimize the number of false positives by identifying variants with characteristics outside their normal distributions: SNPs were excluded if they were supported by reads showing evidence of a strand bias (as estimated by Fisher’s exact test (FS>60.0) or the Symmetric Odds Ratio test (SOR>4.0)) or a bias in the position of alleles within the reads that support them between the reference and alternate alleles (ReadPosRankSum<-8.0). SNPs were also filtered out if they were supported by reads with a low read mapping quality (MQ<40) or a qualitative difference between the mapping qualities of the reads supporting the reference allele and those supporting the alternate allele (MQRankSum<12.5). In addition, SNPs were removed from the data set if the variant confidence was low (QD<2.0). Due to a frequent misalignment of reads in repetitive regions of the genome, leading to an excess of heterozygous genotype calls, SNPs within repeats were excluded from further analyses. In addition, SNPs showing an excess of heterozygosity were removed using the ‘--hardy’ option in VCFtools v.0.1.12b (Danecek *et al.* 2011) with p<0.01. The data set was further limited to SNPs exhibiting complete genotype information.

To achieve a higher specificity, a second independent variant call was performed using Platypus v.0.8.1, an integrated mapping-, assembly-, and haplotype-variant caller (Rimmer *et al.* 2014), and an intersection variant data set was generated.

### Variant Data Set

The intersection data set contained 1,795,643 autosomal variants, with an average transition-transversion (Ts/Tv) ratio of 2.8 (Table 1). Given the use of low coverage (4.5X) sequence data in this study, genotypes were subsequently refined using the software BEAGLE v4 (Browning and Browning 2007). Variants were polarized using rhesus macaque as an outgroup. Using the rhesus macaque genome assembly, rheMac8, consisting of 23 chromosomes as well as 284,705 scaffolds with a total size of 3.2Gb (downloaded from the UCSC Genome Browser), the ancestral state of 1,149,007 variants could be unambiguously determined. For each population, the number of segregating sites shared between any single other population as well as all other populations was recorded, together with the number of segregating sites unique to each population (Figure 2). Subsequently, the data set was annotated using ANNOVAR v2016Feb01 (Wang *et al.* 2010) with the annotation of the vervet genome build (NCBI *Chlorocebus sabaeus* Annotation Release 100) consisting of 29,648 genes, resulting in 22,767 exonic and 577,004 intergenic SNPs.

### Accessible Genome

Due to the fact that the application of filter criteria to the variant data set led to the exclusion of a substantial fraction of genomic sites accessible to variant identification, mask files, defining which genomic sites were accessible to variant discovery, were created. Thereby, monomorphic sites were called and filtered using the same pipeline and hard filter criteria as used for the variant sites (as described in the ‘Variant Calling and Filtering’ section), with the exception of turning the ‘-all’ flag in GATK’s GenotypeGVCFs run on to include non-variant loci. The number of autosomal monomorphic sites in the reference assembly (454,322,622) was then obtained from these mask files. Following the same procedure as for the polymorphic sites, the ancestral state of all monomorphic sites was determined and sites were annotated, resulting in 144,943,664 intergenic monomorphic sites for which the ancestral state could be unambiguously determined.

### Population Structure

A consensus tree *(i.e.,* a tree that most commonly represents the relationship between the sample locations) was constructed based on autosomal variant calls with ancestral allele annotation using the maximum likelihood method implemented in SNPhylo v.20140701 (Lee *et al.* 2014). Gnu R‘s ‘snpgdsLDpruning’ was used with a linkage disequilibrium threshold of r^2^>0.2 to generate a pruned subset of SNPs from the data set, where SNPs are in approximate linkage equilibrium with each other. Using this set of variants, evidence of population structure was assessed using PCA (Zheng *et al.* 2012) as well as an independent-loci admixture model in the software fastSTRUCTURE v.1.0 (Raj *et al.* 2014), detecting clusters of related individuals from multi-locus genotyping data, thereby allowing individuals to have ancestry from multiple populations. fastSTRUCTURE was applied to values of *K* (the number of clusters) between 1 and 5 and the best *K* was chosen such that it maximizes the marginal likelihood. The fraction of identity-by-state (IBS) for each pair of individuals was calculated using Gnu R’s ‘snpgdsIBS’. For each population, summary statistics, namely nucleotide diversity *π* (Nei and Li 1979) and Watterson’s estimate of *θ*, *θ_W_* (Watterson 1975), were calculated using the libsequence package msstats v.0.3.4 (Thornton 2003) (Table 2). Both *π* and *θ_W_* estimate the neutral parameters 4*N_e_μ* under equilibrium conditions, where *N_e_* is the effective population size and *μ* is the neutral mutation rate. Effective population sizes were directly estimated from the data by fixing the mutation rate *μ* to 5.9 × 10^−9^ per base pair per generation (*i.e.,* the mutation rate observed in rhesus macaque, as there is no direct estimate for *μ* available in vervet monkeys; Hernandez *et al.* 2007). Using VCFtools v.0.1.12b (Danecek *et al.* 2011), Tajima’s *D* (Tajima 1989) was calculated (in 10kb windows along the genome) to test for deviations from the equilibrium neutral site frequency distribution, and Weir and Cockerham’s fixation index *F_st_* was calculated between each pair of populations to assess population differentiation.

### Population Divergence

Divergence times were estimated using a molecular clock based on putatively neutral fixed differences (from the intersection data set for which the ancestral state could be unambiguously determined) between the genomes of the populations (Table 3), assuming that mutations occurred at a constant rate *μ* of 5.9 × 10^−9^ per base pair per generation among lineages (Hernandez *et al.* 2007) and a generation time of 8.5 years (Warren *et al.* 2015).

A variety of population bottleneck models were tested and their fit to the data was compared to the fit of a standard equilibrium model. In the proposed bottleneck models, the ancestral effective population size *N_0_* (varied between 10,000, 20,000, 30,000, 40,000, and 50,000 individuals) was reduced to levels of 10%-90% (*i.e.,* the severity of the bottleneck) in 10% intervals for the last 10, 25, 50, 75, and 100 generations (*i.e.,* the duration of the bottleneck). Specifically, for each model, 1,000 independent simulations of 10,000bp length were performed using the coalescence simulator ‘msprime’ (Kelleher *et al.* 2016), assuming a mutation rate *μ* of 5.9 × 10^−9^ per base pair per generation (Hernandez *et al.* 2007) and a recombination rate *ρ*=*μ*. For each simulation, summary statistics (*i.e., F_st_* (Hudson *et al.* 1992) as well as Tajima’s *D* (Tajima 1989)) were calculated using the libsequence package msstats v.0.3.4 (Thornton 2003) and compared to the data. In addition, the fit of a 5-population standard equilibrium model based on the inferred tree topology and the estimated divergence times was assessed.

### Identification of Candidate Loci under Selection

The software SweeD v.3.3.2 (Pavlidis *et al.* 2013), which implements a modification of the Kim and Stephan (2002) composite likelihood ratio (CLR) test as extended by Nielsen *et al.* (2005), was used to detect loci putatively subject to positive selection by scanning the genome for signals of hard (fixed) selective sweeps. For each population, the CLR statistic was calculated from the unfolded SFS at 1,000 grid points across each chromosome. Statistical thresholds were calculated following Nielsen *et al.* (2005) by simulating 1,000 variant data sets under a standard equilibrium model without migration and defining the threshold as the 99^th^ percentile of the distribution of the highest simulated CLR values.

In addition to SweeD, BayeScan v.2.1 (Foll and Gaggiotti 2008) was used to detect loci that show evidence of selection by computing the differences in allele frequencies between the different populations.

## Acknowledgements

I am grateful to Jeffrey Jensen and Stefan Laurent for helpful comments and discussion. Computations were performed at the Vital-IT (http://www.vital-it.ch) Center for high-performance computing of the Swiss Institute of Bioinformatics (SIB).

